# Large-Scale Brain Networks Track Gasoline Price Shifts

**DOI:** 10.64898/2025.12.16.694768

**Authors:** Muwei Li

**Affiliations:** Vanderbilt University Institute of Imaging Science, Vanderbilt University Medical Center, Nashville, TN, USA; Department of Radiology and Radiological Sciences, Vanderbilt University Medical Center, Nashville, TN, USA

## Abstract

Macroeconomic conditions shape daily constraints and perceived environment, yet it remains unclear whether real-world economic dynamics are reflected in intrinsic brain organization at the population level. Here, I leveraged the Human Connectome Project (HCP) acquisition timeline as a naturalistic sampling frame to test whether macroeconomic time series vary with large-scale resting-state networks. Resting-state fMRI from 726 healthy young adults was aligned to quarterly macroeconomic indicators using the HCP “Quarter” variable. I quantified intrinsic organization using network functional connectivity (FC) and network-level amplitude of low-frequency fluctuations (ALFF). I implemented (1) a quasi-natural experiment contrasting a pre- vs post-gasoline-price shock (collapse) cohort, and (2) quarter-level partial correlations between network measures and four macro indicators, including gasoline price, consumer sentiment, unemployment, and stock market return. Post-shock participants showed significantly higher within-network FC across all seven networks and selective increases in between-network coupling concentrated among sensory and attention systems, as well as limbic-default interactions. ALFF exhibited bidirectional shifts: Visual ALFF increased post-shock, whereas Limbic and Frontoparietal ALFF were higher pre-shock. Quarter-level analyses mirrored this differential pattern, with gas price positively associated with Limbic and Frontoparietal ALFF and negatively with Visual and Dorsal Attention ALFF. Across macro indicators, gasoline price and consumer sentiment showed the most widespread FC associations, with largely opposing correlation signatures, while unemployment and equity returns were comparatively weak after correction. These findings suggest that salient, behaviorally proximal macroeconomic dynamics, particularly energy price variation, track population-level differences in intrinsic brain network architecture, motivating future work with finer-grained timing and individual-level exposure measures to strengthen causal inference.

## INTRODUCTION

Macroeconomic conditions are not merely abstract background variables. They shape the day-to-day constraints under which people plan, travel, spend, and cope. Large downturns can have persistent effects on mental health, especially for individuals who experience direct financial, job, or housing losses (Forbes and Krueger 2019). Across longitudinal evidence, unemployment is associated with increases in depression, anxiety, and psychological distress, and symptom levels tend to improve after re-employment (Sterud et al. 2025). Conversely, quasi-random positive income shocks, such as medium-sized lottery wins, predict later improvements in psychological well-being (Gardner and Oswald 2007). These population-level observations align with an allostasis perspective in which the brain dynamically adapts to changing environmental demands, while repeated adaptation can accumulate “wear and tear” that influences health and behavior (Ganzel et al. 2010). If macroeconomic shifts measurably alter stress exposure, affect, and everyday choices, they may also be reflected in the organization of human brain function at the population level.

Neuroimaging research has established multiple links between socioeconomic conditions and the brain, spanning both structural and functional domains. Structurally, neuroimaging studies have linked socioeconomic context to variation in brain regions implicated in memory, emotion regulation, language, and executive control. These associations are often discussed in relation to material deprivation, caregiving environments, stress exposure, and environmental toxins as plausible mediating pathways (Brito and Noble 2014; Johnson et al. 2016; Olson et al. 2021). Specific exposures such as current financial hardship have been associated with smaller hippocampal and amygdala volumes (Butterworth et al. 2012), and cumulative physiological burden (allostatic load) has been linked to brain volume and cognitive ability in aging (Booth et al. 2015) as well as cortical thickness patterns in adulthood (Ottino-González et al. 2017). Diffusion MRI studies further show socioeconomic gradients in white-matter integrity (Johnson et al. 2013; Shaked et al. 2019), which may partially mediate socioeconomic disparities in executive function (Shaked et al. 2022). Longer-horizon economic instability also matters. Income volatility across early-to-mid adulthood has been associated with poorer cognition and reduced white matter integrity in midlife (Grasset et al. 2019), and life-course socioeconomic conditions have been linked to quantitative markers of white matter microstructure and cognitive performance (Schrempft et al. 2024).

Functional studies have approached the socioeconomic-brain question through both electrophysiology and task fMRI. In early development, randomized cash-transfer studies demonstrated that poverty-alleviation participation has been linked to lower baseline cortisol in children (Fernald and Gunnar 2009), and unconditional cash gifts have been associated with shifts in resting EEG power measured from infants (Troller-Renfree et al. 2022). Some other EEG studies report socioeconomic associations with resting oscillatory activity and developmental trajectories, spanning infancy through school age (Maguire and Schneider 2019; Jensen et al. 2021; Sandre et al. 2025), while ERP work suggests socioeconomic differences in attentional control and related neuroendocrine correlates (D’angiulli et al. 2012). Task fMRI studies often frame economic constraints through reward valuation and cognitive control systems, suggesting that subjective value is represented in a distributed network including ventral striatum and medial prefrontal cortex (Kable and Glimcher 2007), and prefrontal effective connectivity helps explain individual differences in temporal discounting during monetary choice (Hare et al. 2014). Importantly, experimentally induced scarcity mindsets can shift these computations, enhancing orbitofrontal valuation signals while reducing dorsolateral prefrontal control activity (Huijsmans et al. 2019), and perceived scarcity can impair cognitive flexibility with identifiable electrophysiological signatures (Huang et al. 2023). Moreover, scarcity manipulations may also modulate socioaffective processing, including empathy-related neural responses (Li et al. 2025).

Together, this literature indicates that both chronic socioeconomic conditions and acute perceptions of scarcity can influence neural systems supporting attention, valuation, and control.

Despite these advances, two gaps limit our understanding of how broad economic environments relate to brain function. First, most prior neuroimaging work measures exposure at the individual or household level (e.g., SES, poverty, financial hardship), while macroeconomic time-varying indicators, such as energy prices, consumer sentiment, unemployment, or market volatility, are rarely used as explicit exposures in neuroimaging models. Second, much of the functional literature has relied on task paradigms that operationalize economic influences through reward valuation and cognitive control manipulations, rather than leveraging real-world macroeconomic time series to probe intrinsic network organization. While valuable, this emphasis leaves open a complementary question: whether real-world macroeconomic dynamics, shared across large populations and fluctuating over time, are reflected in the intrinsic organization of large-scale brain networks (i.e., within- and between-network coupling) and the degree of brain activity rather than only in task-evoked contrasts.

I address these gaps by leveraging a natural experiment embedded in the acquisition timeline of the Human Connectome Project (HCP) (Van Essen et al. 2013; Elam et al. 2021). HCP resting-state fMRI data were collected across multiple quarters, providing a unique opportunity to align each participant with macroeconomic conditions at the time of scanning and to examine population-level brain organization as a function of macroeconomic variation. I focus on gasoline price dynamics for three reasons. First, during HCP data collection, a major gasoline price collapse occurred, providing an externally defined regime shift that allows a complementary “pre/post shock” analysis in addition to continuous across-quarter modeling. Second, gasoline prices are unusually salient and behaviorally proximal. They directly affect commuting, transportation costs, and household budgets, making them a widely perceived component of everyday economic strain. Third, evidence from economics demonstrates that gasoline prices can shape durable behavior. Exposure during formative years predicts reduced driving decades later, highlighting gasoline price as a meaningful population-level signal of perceived cost and constraint (Severen and van Benthem 2022). These considerations motivate testing whether gasoline prices, benchmarked against other economic indicators, vary with intrinsic brain network organization.

In this study, I quantify intrinsic brain organization using two complementary measures: (1) within- and between-network functional connectivity (FC) across canonical large-scale networks (Yeo networks), and (2) amplitude of low-frequency fluctuations (ALFF) as a network-level index of the magnitudes of BOLD dynamics. First, I treat the gasoline price collapse as a quasi-natural experiment and compare pre-versus post-shock cohorts in within- and between-network FC and in network-level ALFF. Second, I model macro-brain coupling continuously by correlating quarterly network FC and ALFF with multiple macroeconomic indicators (including gasoline price and comparator variables such as consumer sentiment, unemployment, and equity market returns). Across analyses, I test whether gasoline prices show the strongest and most consistent associations with intrinsic network organization. By integrating a discrete shock-based comparison with a continuous across-quarter framework, my results suggest that macroeconomic dynamics, particularly energy price variation, are reflected in large-scale intrinsic brain networks, providing a scalable paradigm for studying how societally shared environmental fluctuations relate to population-level brain function.

## MATERIALS AND METHODS

### Ethics Statement

The data involved in this research are publicly available and have been previously approved for use by the Washington University Institutional Review Board. All participants provided written informed consent to participate in this study. The authors did not collect any new data involving human participants.

### Economic Data

I examined four U.S. macroeconomic indicators that were available at regular (monthly or higher) frequency over the HCP acquisition window. (1) Gasoline price (weekly, U.S. regular retail). Retail gasoline prices were obtained from the U.S. Energy Information Administration (eia.gov) historical data series (Weekly U.S. Regular All Formulations Retail Gasoline Prices; dollars per gallon). (2) S&P 500 (stock market index) total return (monthly, including dividends). Monthly S&P 500 total returns were obtained from OfficialData.org, which reports returns derived from the Robert Shiller S&P 500 dataset. (3) U.S. unemployment rate (monthly, seasonally adjusted). Monthly unemployment rates were obtained from the Federal Reserve Bank of St. Louis FRED database (fred.stlouisfed.org), defined as the number of unemployed persons as a percentage of the labor force. (4) Consumer sentiment (monthly). Monthly consumer sentiment was obtained from the University of Michigan Consumer Sentiment Index accessed via FRED (fred.stlouisfed.org), with the original source being the University of Michigan Surveys of Consumers. All macroeconomic series were aligned to the HCP acquisition-quarter index by averaging the underlying time series within each quarter window.

### Imaging Data

Because the Human Connectome Project Young Adult data (Van Essen et al. 2013) does not publicly release exact scan dates for individual participants (for privacy and data-governance reasons), I used the HCP-provided Quarter variable as the temporal index for aligning each participant to macroeconomic conditions. This variable indicates the quarter in which 3T imaging and behavioral data of the subjects were initially acquired. In total, it comprises 13 quarters: Q1 = Aug-Oct 2012; Q2 = Nov 2012-Jan 2013; Q3 = Feb-Apr 2013; Q4 = May-Jul 2013; Q5 = Aug-Oct 2013; Q6 = Nov 2013-Jan 2014; Q7 = Feb-Apr 2014; Q8 = May-Jul 2014; Q9 = Aug-Oct 2014; Q10 = Nov 2014-Jan 2015; Q11 = Feb-Apr 2015; Q12 = May-Jul 2015; and Q13 = Aug-Oct 2015. I excluded Q1-Q3, as these quarters used an image reconstruction approach that was not fully consistent with later data releases. Therefore, from Q4-Q13, I selected 726 entries (comprising 360 males and 366 females, all between the ages of 22 and 35 years), adhering to criteria that included the completeness of a 3T scan, along with adequate data quality.

I defined a gasoline-price “shock” around the 2014 collapse in crude oil prices. The U.S. Energy Information Administration (https://www.eia.gov/finance/review/annual/) summarizes that Brent prices peaked on 2014-06-19 and that the second half of 2014 saw a pronounced decline, with major drivers including rising global supply, fewer supply disruptions, downward revisions to demand growth expectations, and OPEC maintaining production levels. I therefore categorized participants into two groups: Pre-shock (Q4-Q9, May 2013-Oct 2014) and Post-shock (Q10-Q13, Nov 2014-Oct 2015). In my sample, the pre-shock group comprised 467subjects (F = 244, M=223, age: 22-35), and the post-shock group comprised 259 subjects (F = 122, M = 137, age: 22-35).

The imaging protocols are detailed elsewhere (Van Essen et al. 2012) and were performed with 3T Siemens Skyra scanners. Two sessions of resting-state scans were acquired using multiband gradient-echo EPI sequences. Each session consisted of two runs of scans with opposing phase encoding directions and lasted 14 minutes and 33 seconds, with parameters TR = 720 ms, TE = 33.1 ms, and an isotropic voxel resolution of 2 mm, totaling 1200 volumes. Additionally, T1-weighted images were obtained using a single-echo MPRAGE sequence with a TR of 2,400 ms, TE of 2.14 ms, and voxel dimensions of 0.7 mm isotropic.

### Preprocessing

T1-weighted images are nonlinearly registered to MNI space using FNIRT (Jenkinson et al. 2012), and cortical surface reconstructions are generated using FreeSurfer (Dale et al. 1999). These surfaces were then registered to a standard template using MSMAll (Robinson et al. 2018), which aligns cortical areas based on multimodal features rather than geometry alone. Functional preprocessing includes motion correction, distortion correction (Andersson et al. 2003), and registration to the corresponding structural image, followed by nonlinear alignment to MNI space. After spatial normalization, the data were projected to the standard 32k fs LR surface mesh and saved in CIFTI dtseries. Additional preprocessing includes denoising via ICA-FIX (Salimi-Khorshidi et al. 2014), which removes structured noise components while preserving the neural signal, and nuisance regression of motion parameters. I further performed linear detrending and band-pass filtering (0.01-0.1 Hz). After preprocessing, regional time series were extracted using the HCP-MMP1.0 multimodal parcellation (Glasser et al. 2016), which defines 360 cortical parcels (180 per hemisphere) aligned to the same 32k fs LR surface space as the functional data. The atlas was applied directly to the CIFTI dtseries files using HCP workbench software. For each participant and each run, this procedure yielded a 360 × T matrix of parcel-averaged BOLD time series, where T corresponds to the number of volumes in the run (T = 1200).

### FC and ALFF Calculation

Resting-state FC was computed from the parcel-wise time series described above. For each participant, I used all four resting-state runs. Within each run, pairwise FC between all parcel pairs was estimated using the Pearson correlation coefficient, yielding a 360 × 360 correlation matrix R, and each element was then transformed to Fisher-z values. Fisher-z FC matrices were subsequently averaged across runs to obtain a single participant-level 360 × 360 Fisher-z connectivity matrix. Each of the 360 parcels was assigned to one of the seven canonical Yeo networks (Thomas Yeo et al. 2011) (Visual, Somatomotor, Dorsal Attention, Ventral Attention, Limbic, Frontoparietal, Default). The participant-level 360 × 360 Fisher-z FC matrix was then reduced to a 7 × 7 network-level matrix by averaging Fisher-z values within and between Yeo networks. Specifically, each diagonal entry (within-network FC) was computed as the mean Fisher-z across all unique parcel pairs within the same network (upper triangle of the corresponding block, excluding the diagonal). Each off-diagonal entry (between-network FC) was computed as the mean Fisher-z across all parcel pairs spanning the two networks (i.e., all elements in the corresponding off-diagonal block).

ALFF was calculated from the parcel-wise resting-state time series for each participant using four resting-state runs. For each run and each parcel, I first removed the mean from the parcel time series and then computed the discrete Fourier transform (FFT). The single-run ALFF for each parcel was defined as the mean amplitude of the Fourier magnitude spectrum within the 0.01-0.10 Hz band. Run-wise ALFF estimates were then averaged across runs to yield a single participant-level ALFF vector (360 ×1). I additionally computed a z-scored ALFF (zALFF) by standardizing the 360 parcel ALFF values within participant (subtracting the across-parcel mean and dividing by the across-parcel standard deviation). Finally, parcel-level zALFF was summarized to the Yeo-7 network level by averaging ALFF values across parcels assigned to each network (Visual, Somatomotor, Dorsal Attention, Ventral Attention, Limbic, Frontoparietal, Default).

### Statistical Analyses

Statistical analyses were performed at the Yeo-7 network level (7 within-network measures and 21 unique between-network pairs; 28 total features). To test pre-shock vs post-shock differences in network FC, I fit multiple linear regression models with group (pre/post) as the predictor of interest and head motion and gender as covariates. Significance was assessed on the group term and FDR-corrected across all 28 network edges (q < 0.05). For network ALFF, pre/post comparisons were performed within each network and corrected using FDR across the seven networks. To relate brain measures to quarter-level macroeconomic indicators (GasPrice_q, SP500_ret_q, UnempRate_q, UMichSent_q), I computed partial correlations (r) between each network-level feature and each macroeconomic series while controlling for motion and gender, and applied FDR correction across the 28 network features for each macroeconomic variable. Finally, for ALFF-macroeconomic associations at the cortical level, I computed parcel-wise partial correlation maps (controlling for motion and gender) and summarized effects as network-wise partial correlations, with FDR correction across the seven networks for each macroeconomic variable.

## RESULTS

### Network FC Shifts After the Shock

Figure 1 summarizes how large-scale network FC differed between the pre-shock (high price) and post-shock (low price) quarters. At the group level (Fig. 1a), the canonical Yeo-7 architecture was preserved in both regimes, but the post-pre difference matrix showed a broad upward shift in connectivity, with multiple edges surviving FDR correction. Consistent with this pattern, within-network FC was significantly higher in the post-shock group across all seven networks (Fig. 1b; FDR-corrected q-values indicated by asterisks). Between-network effects were more selective (Fig. 1c). Significant post-shock increases were concentrated among sensory/attention systems (e.g., Visual-Somatomotor and Visual-Attention links; Somatomotor-Attention links; and Dorsal-Ventral Attention coupling), with additional effects involving Limbic interactions (including attention-limbic coupling and limbic-default coupling), whereas many other cross-network pairs were not significant after FDR correction.

**Figure 1.**
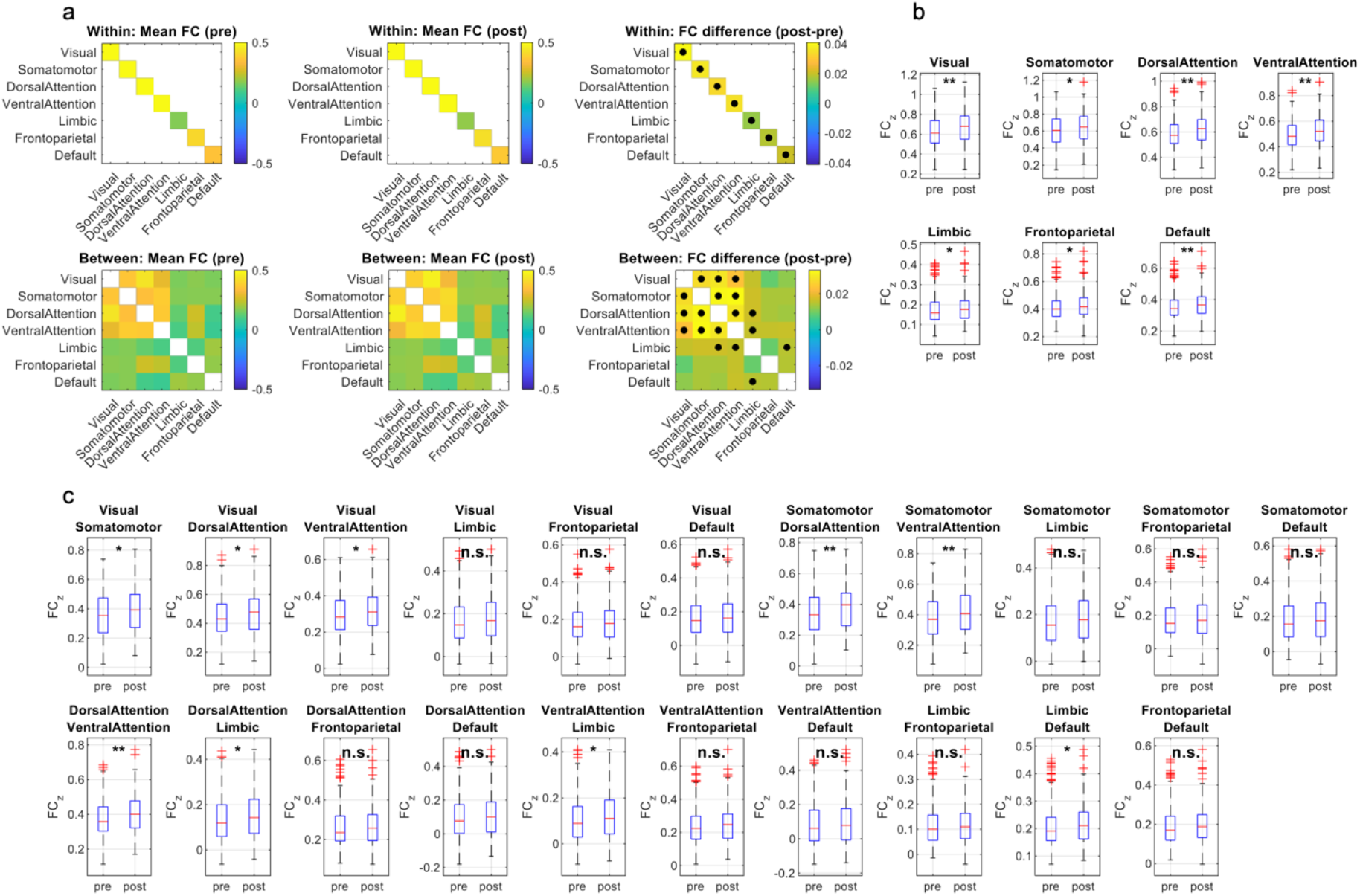
Network-level functional connectivity across high- vs low-gasoline price regimes. (a) Group-mean FC among the seven Yeo networks for pre-shock and post-shock subjects, shown separately for within-network (top row) and between-network (bottom row) connections, together with post minus pre difference matrices. Black dots mark connections whose FC shows a significant effect of pre vs post in a multiple linear regression controlling for head motion and gender, FDR-corrected across all 7 within- and 21 between-network edges (q < 0.05). (b) Boxplots of within-network FC for each Yeo network, comparing pre-shock (pre) and post-shock (post) quarters. (c) Boxplots of between-network FC for all 21 unique network pairs, ordered consistently with panel (a). Asterisks on top of the boxplots indicate the FDR-corrected significance (* q < 0.05, ** q < 0.01; “n.s.”, not significant).

### Network ALFF Shows Bidirectional Shifts After the Shock

Figure 2 shows that the gasoline-price regime was associated with selective shifts in intrinsic ALFF rather than a uniform change across networks. In the pre/post comparison (Fig. 2a), Visual ALFF was significantly higher in the post-shock (low-price) group, whereas Limbic and Frontoparietal ALFF were significantly higher in the pre-shock (high-price) group (FDR-corrected across the seven networks; other networks were not significant). Consistent with this pattern, quarter-level analyses (Fig. 2b) revealed significant partial correlations between network ALFF and the quarter-level gasoline-price index (GasPrice_q), controlling for motion and gender: Limbic and Frontoparietal networks showed positive associations (higher ALFF during more expensive quarters), while Visual and Dorsal Attention networks showed negative associations (lower ALFF during more expensive quarters), surviving FDR correction across networks.

**Figure 2.**
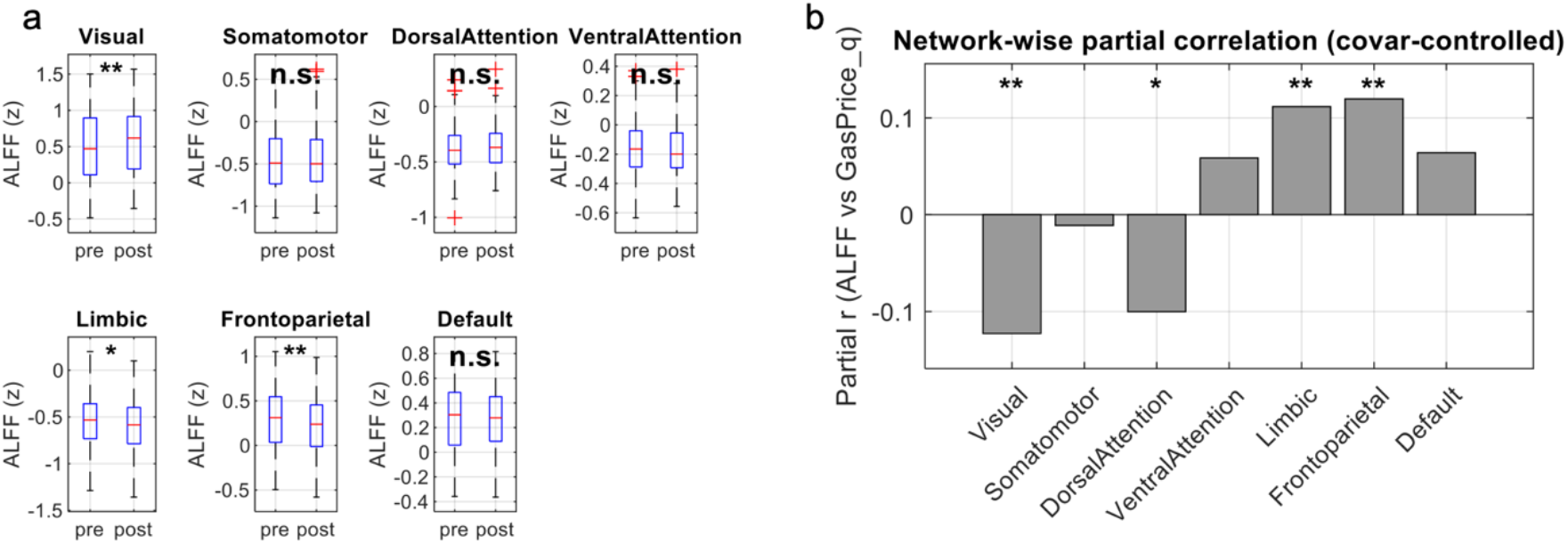
Network-level ALFF across high- vs low-gasoline price regimes. (a) Boxplots show z-scored ALFF (0.01-0.10 Hz) within each Yeo-7 network for subjects scanned in high-price “pre-shock” quarters versus low-price “post-shock” quarters. Asterisks above the boxes indicate the FDR-corrected significance of the comparisons (* q < 0.05, ** q < 0.01; n.s., not significant). (b) Bars plot the partial correlation between ALFF and the gasoline-price index GasPrice_q, defined as the mean national gasoline price in the acquisition quarter of a subject, controlling for motion and gender. Positive values indicate higher ALFF during more expensive quarters. Asterisks denote networks with a significant correlation after FDR correction across networks (* q < 0.05, ** q < 0.01).

### Correlation between Macroeconomic and FC

Figure 3 extends the gasoline-price findings by evaluating three additional macroeconomic indicators primarily as complementary controls. Overall, the strongest and most widespread associations with network FC were observed for gasoline price and consumer sentiment, whereas S&P 500 returns and unemployment showed comparatively weak or nonsignificant relationships at the network level after multiple-comparison correction. At the within-network level (Fig. 3a-d), gasoline price (GasPrice_q) was significantly related to within-network FC across multiple networks, with the direction of effects varying by system. In contrast, consumer sentiment (UMichSent_q) exhibited broadly positive within-network associations that survived FDR correction across networks. At the between-network level (Fig. 3e-h), gasoline price showed a broad pattern of significant associations across many cross-network pairs (predominantly negative), indicating that higher gasoline prices were associated with reduced integration between large-scale systems. Consumer sentiment again demonstrated widespread significant positive associations across between-network pairs, consistent with a global upshift in cross-network coupling during quarters with higher sentiment. Black markers denote effects surviving FDR correction across the 28 network features for each macroeconomic variable (q < 0.05).

**Figure 3.**
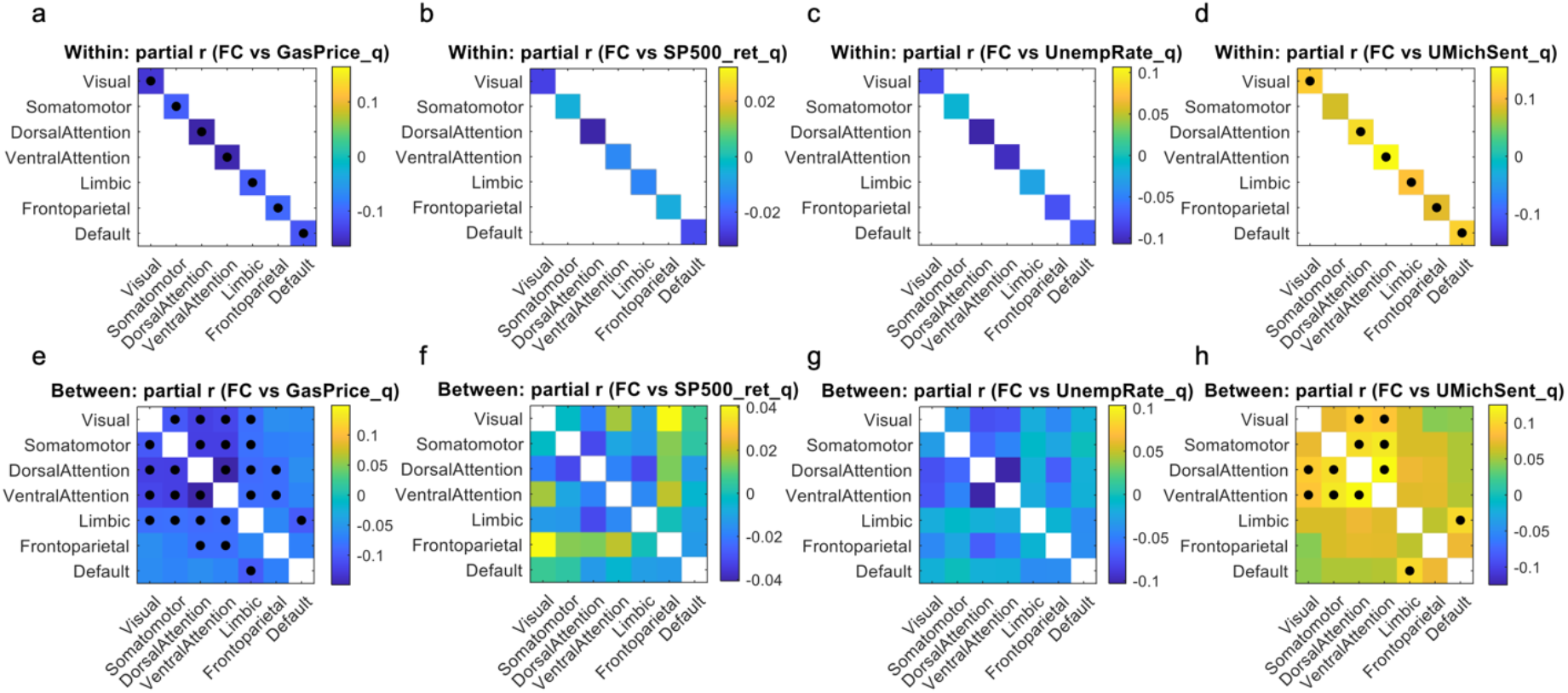
Correlations between large-scale network FC and quarter-level macroeconomic indicators. Panels a-d show within-network effects (only diagonal entries are available), and panels e-h show between-network effects (only off-diagonal entries). Color indicates the partial correlation coefficient r between FC and: (a,e) mean U.S. gasoline price (GasPrice_q), (b,f) S&P 500 total return (SP500_ret_q), (c,g) national unemployment rate (UnempRate_q), and (d,h) University of Michigan Consumer Sentiment (UMichSent_q), all defined at the HCP-quarter level. Black dots mark connections with a significant association after FDR correction across all 28 network features for that economic variable (q < 0.05).

### Macroeconomic Context for ALFF Findings

Figure 4 summarizes how resting-state ALFF varies with quarterly macroeconomic indicators after covariate control. Among the four macro indicators, gasoline price showed the clearest and most robust ALFF associations (Fig. 4a-b). At the network level, higher gas prices were linked to lower ALFF in the Visual network (largest negative effect, FDR-corrected) and in Dorsal Attention (negative, FDR-corrected), but higher ALFF in Limbic and Frontoparietal networks (both positive; FDR-significant), with smaller and non-significant effects in the remaining networks. The cortical map echoed this pattern, with cooler colors in occipital/visual territories and warmer colors in higher-order association cortex. In contrast, ALFF associations with the other three indicators (S&P 500 return, unemployment rate, and consumer sentiment) were comparatively modest at the network level (Fig. 4d, f, h) and did not show FDR-surviving network effects, although their spatial maps suggested weaker, distributed trends (e.g., consumer sentiment exhibiting a largely opposite sign pattern relative to gas price, with more positive effects in Visual/Dorsal Attention and negative tendencies in Limbic/Frontoparietal; Fig. 4g-h).

**Figure 4.**
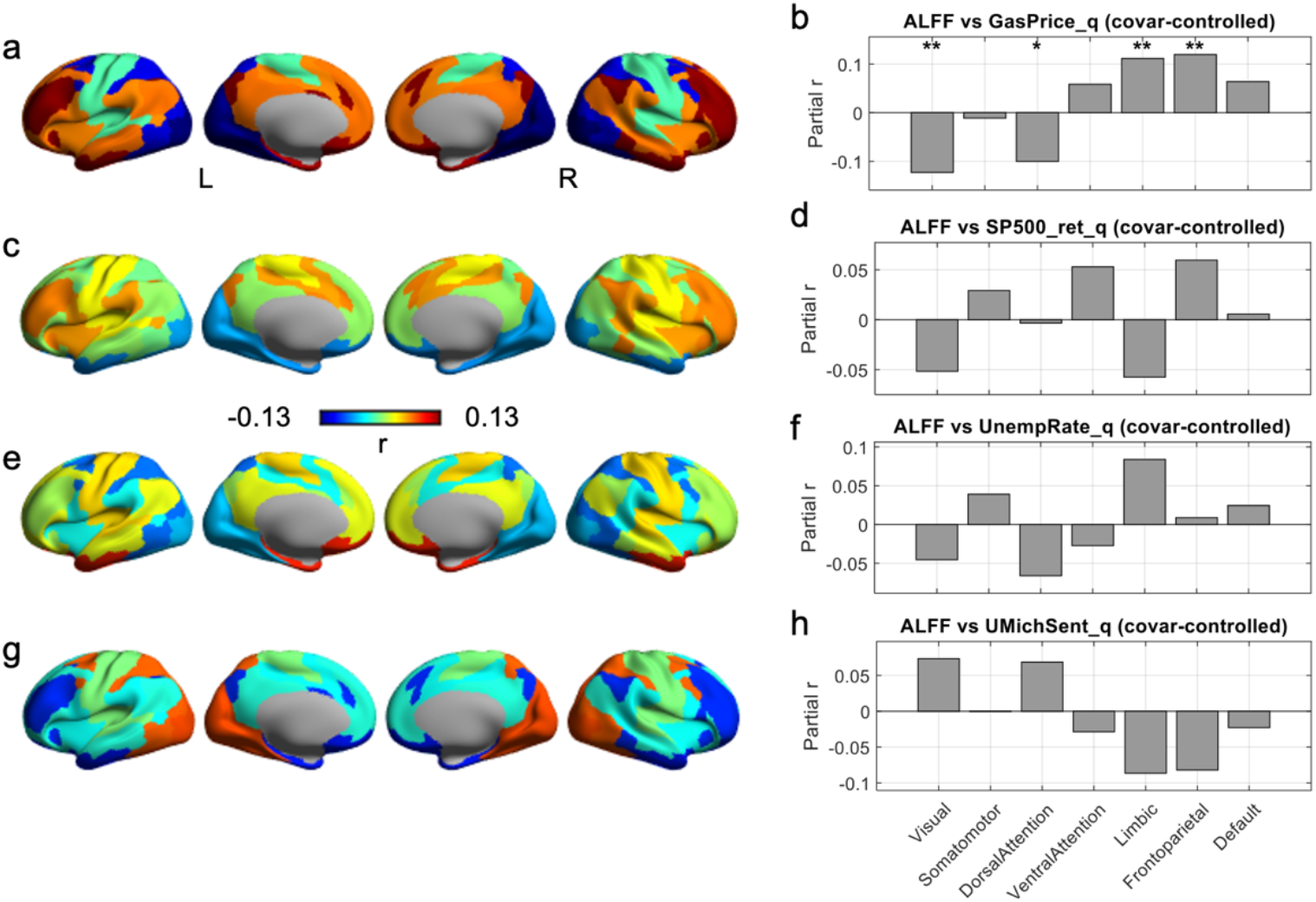
Cortical distribution of ALFF-macroeconomic associations. (a-b) Gasoline price. Panel a shows the cortical map of partial r between ALFF and quarter-level gasoline price (GasPrice_q); warm colors indicate higher ALFF when gas prices are higher, cool colors indicate the opposite. Panel b shows the corresponding network-wise bar plot. (c-d) S&P 500 returns. Panel c shows surface maps of partial r between ALFF and quarterly S&P 500 total return (SP500_ret_q); panel d gives the corresponding network-wise bar plot. (e-f) Unemployment rate. Panel e displays partial r between ALFF and the national unemployment rate (UnempRate_q); panel f shows the network-wise values. (g-h) Consumer sentiment. Panel g shows partial r between ALFF and the University of Michigan Consumer Sentiment index (UMichSent_q); panel h summarizes the seven network values. In all bar plots (b,d,f,h), gray bars give the partial correlation coefficient for each network, controlling for motion and gender. Asterisks mark networks that survive correction across the seven networks for that macro variable (* q < 0.05, ** q < 0.01; n.s., non-significant effects).

## DISCUSSIONS

This study tested whether real-world macroeconomic dynamics-treated as societally shared, time-varying exposures, are reflected in the intrinsic organization of large-scale brain networks. Leveraging the Human Connectome Project acquisition timeline as a naturalistic sampling frame, I aligned each participant to quarterly windows and examined resting-state network FC and low-frequency BOLD amplitude in relation to a major gasoline-price regime shift and to quarter-level macroeconomic indicators. Three findings stand out. First, the gasoline price collapse provided a clear “pre/post” contrast in which post-shock participants showed a broad increase in within-network FC across all seven canonical networks, with more selective increases in between-network coupling concentrated among sensory and attention systems and their interactions with limbic/default networks. Second, ALFF did not show a uniform shift but instead exhibited bidirectional, network-specific changes: Visual ALFF was higher post-shock, whereas Limbic and Frontoparietal ALFF were higher pre-shock. Third, when comparing gasoline price with three complementary macroeconomic series (consumer sentiment, unemployment, and S&P 500 returns), gasoline price and consumer sentiment showed the most robust and widespread associations with network FC, whereas the other indicators were comparatively weak after correction. Together, these results support the central premise that macroeconomic time series that are salient in everyday life vary with large-scale intrinsic brain network organization at the population level.

The FC findings suggest that macroeconomic regimes can be accompanied by coherent, system-level shifts in intrinsic coupling. The most consistent effect was the elevation of within-network FC across all Yeo networks in the post-shock cohort. Because within-network FC indexes the degree of coordinated activity among parcels belonging to the same functional system, a global post-shock increase is consistent with a broad change in network “cohesion” or internal synchrony. Importantly, this effect was observed across networks with diverse functional roles (sensory, attention, control, limbic, default), indicating that the shift is unlikely to reflect a single domain-specific process (e.g., only reward-related circuitry). Instead, it may reflect a general change in the intrinsic organizational state under different macroeconomic conditions. Between-network effects were more selective, with significant post-shock increases concentrated among sensory and attention systems (Visual-Somatomotor, Visual-Attention, Somatomotor-Attention, Dorsal-Ventral Attention) and with additional effects involving limbic interactions (e.g., attention-limbic coupling and limbic-default coupling). One interpretation is that macroeconomic context may differentially influence the balance between segregated and integrated processing, with some cross-system communication channels being more sensitive than others. Sensory and attentional systems are tightly linked to everyday behavior, i.e., how people allocate attention, plan actions, and engage with the external world (Land and Hayhoe 2001; Shamay-Tsoory and Mendelsohn 2019), and may therefore be among the first systems to show population-level signatures of shifting constraints and routines. The fact that the limbic and default-mode networks also changed suggests that the macroeconomic context may influence not only attention/sensory processing, but also mood-related and internally focused processes (e.g., emotion, self-referential thought, mind-wandering). This fits an allostasis view: the brain continually adjusts its activity to meet changing environmental demands, and if these adjustments are sustained over time, they may come with a cumulative ‘wear-and-tear’ cost on the body and mind (McEwen and Gianaros 2011; Kleckner et al. 2017).

In contrast to FC, ALFF revealed a dissociative pattern that is difficult to explain as a single global arousal effect. Visual ALFF increased in the post-shock period, whereas Limbic and Frontoparietal ALFF were higher in the pre-shock period, and quarter-level correlations with gasoline price showed the same directionality (positive for Limbic/Frontoparietal, negative for Visual/Dorsal Attention). This bidirectional organization suggests a redistribution of low-frequency intrinsic dynamics across systems rather than a uniform shift. Under relatively higher gasoline prices (interpretable as a more salient cost environment), increased Limbic and Frontoparietal ALFF could reflect greater internally maintained affective load or increased reliance on control-related systems involved in coping, planning, and regulation (Haushofer and Fehr 2014; Huijsmans et al. 2019). Conversely, reduced ALFF in the Visual and Dorsal Attention networks during higher-price quarters may indicate attenuated spontaneous low-frequency dynamics in externally oriented processing systems, a pattern that is broadly consistent with the FC results suggesting altered network-level engagement with the external environment. While these interpretations remain speculative, the key point is that the macroeconomic signal is expressed as a structured network-level “reorganizing” that aligns with the distinction between sensory/attention systems and higher-order limbic/control systems.

Across macroeconomic indicators, gasoline price and consumer sentiment exhibited the strongest and most widespread associations with network FC, whereas unemployment and equity market returns were comparatively weak after multiple-comparison correction. One important consideration, however, is that the HCP acquisition window coincided with substantially greater variability in gasoline prices than in the other indicators, which could increase statistical power and partially contribute to the prominence of gas-related effects. Even with this caveat, the pattern is still informative because it argues against a trivial explanation in which any macro series correlates with brain measures via shared time trends. Instead, it suggests that specific macroeconomic signals, particularly those that are behaviorally proximal or psychologically salient, may be more tightly coupled to intrinsic network organization. Gasoline price is highly visible and frequently encountered through commuting and household budgets and thus may plausibly influence perceived constraints and daily decisions even among individuals not directly engaged with financial markets. Consumer sentiment, in turn, can be viewed as an aggregate index of expectations and affective appraisal of economic conditions. Notably, in my data, gasoline price and consumer sentiment showed largely opposing correlation signatures across networks and edges, providing an internal reproducibility check. This opposition also supports the interpretation that distinct components of “economic context” may be reflected in brain organization, one capturing concrete cost pressure (gas price) and another capturing broader optimism (sentiment). Future work should explicitly model shared versus unique variance across indicators (e.g., multivariable) and quantify how much of the gas advantage is explained by its higher volatility versus its psychological proximity.

Most socioeconomic neuroimaging studies emphasize individual-level exposures (SES, poverty, hardship) or relatively static structural outcomes. Functional studies often operationalize “economics” through laboratory tasks manipulating reward, value, or impulsivity. My findings complement these lines of work by demonstrating that macroeconomic time series-shared at the population level and conceptually, this extends the socioeconomic-brain literature from stable person-level traits and task-evoked circuits to a naturalistic, time-varying exposure framework in which the macro environment serves as an implicit contextual input. In this view, intrinsic network coupling and low-frequency dynamics provide a potential population-level readout of how broad societal conditions may be reflected in brain organization.

Several limitations temper interpretation. First, HCP does not provide exact scan dates. Quarter windows are coarse temporal bins. Aligning macroeconomic series by averaging within each quarter introduces temporal imprecision and may attenuate effects. The fact that robust associations were still observed for gasoline price suggests that the coupling may be relatively strong, but the temporal coarseness prevents testing finer-grained timing hypotheses (e.g., lagged effects). Second, the design is not longitudinal within individuals. It compares different participants scanned in different quarters. Therefore, the results do not establish that macroeconomic changes caused within-person changes in brain organization. Third, macroeconomic indicators were necessarily treated as national-level signals, whereas individuals differ widely in their exposure and sensitivity to gasoline prices (e.g., commuting distance, income, urban versus rural living). The absence of individual-level economic exposure measures limits mechanistic claims. Finally, generalizability is constrained by the HCP-Y sample (healthy young adults). Macroeconomic stressors may exert stronger or qualitatively different effects in more vulnerable groups (adolescents, older adults, individuals with psychiatric risk, or economically disadvantaged populations).

In sum, by combining a shock-based pre/post comparison with continuous quarter-level macro-brain coupling, this study provides evidence that macroeconomic dynamics-especially gasoline price variation-are reflected in the intrinsic functional architecture of the human brain. The observed pattern involves both broad changes in within-network coupling and structured, network-specific shifts in low-frequency dynamics. These results support a scalable paradigm for studying how societally shared environmental fluctuations relate to population-level brain function and open new avenues for connecting macroeconomic context to intrinsic network organization beyond traditional individual-level SES and task-based approaches.

## ACKNOWLEDGMENTS

I am grateful to Dr. John Gore for his continuous support, mentorship, and insightful discussions throughout this work. I also thank Dr. Zhaohua Ding for valuable feedback on the analyses and interpretation of the results.

Imaging data were provided by the Human Connectome Project, WU-Minn Consortium (Principal Investigators: David Van Essen and Kamil Ugurbil; 1U54MH091657), funded by the 16 NIH Institutes and Centers that support the NIH Blueprint for Neuroscience Research; and by the McDonnell Center for Systems Neuroscience at Washington University.

## AUTHOR CONTRIBUTIONS

Muwei Li (Conceptualization, Formal analysis, Investigation, Methodology, Software, Validation, Visualization, Writing)

## Conflict of interest statement

None declared.

## Notes

### Competing Interest Statement

The authors have declared no competing interest.

